# Ancient *Yersinia pestis* genomes provide no evidence for the origins or spread of the Justinianic Plague

**DOI:** 10.1101/819698

**Authors:** Marcel Keller, Maria A. Spyrou, Michael McCormick, Kirsten I. Bos, Alexander Herbig, Johannes Krause

## Abstract

Along with the publication of 137 ancient human genomes retrieved from archaeological remains of the Eurasian steppe, Damgaard et al., 2018 identified two individuals infected with *Yersinia pestis*, yielding one genome with 0.24x average coverage (DA147, 6^th^–9^th^ c. AD) and another with 8.7x (DA101, 2^nd^–3^rd^ c. AD). A phylogenetic analysis performed on the latter placed it in a position ancestral to a 6^th^-century Justinianic genome from Aschheim, Germany. These results are used to fuel an argument that the Justinianic Plague (541–544 AD) “was brought to Europe towards the end of the Hunnic period through the Silk Road along the southern fringes of the steppes” in contrast to the leading hypothesis of introduction via the Red Sea that is supported by historical accounts. In our reanalysis, we question the contested historical context of the presented genomes with the Justinianic Plague and show that the lower coverage genome might be rather related to the Black Death (1346–1353 AD).

## Introduction

The recent sequencing of dozens of pathogen genomes reconstructed from ancient DNA enabled increased-resolution phylogeographic studies on the spread of infectious diseases in prehistoric and historic times, especially in the context of human migration, mobility and trade (Andrades Valtueña et al., 2017; Bos et al., 2016; Keller et al., 2019; Namouchi et al., 2018; Rascovan et al., 2019; Rasmussen et al., 2015; Spyrou et al., 2019, 2016; Vågene et al., 2018). This is especially true for plague with its long and richly documented history and the abundance of published ancient genomes of its causative agent, *Yersinia pestis*. Interpretation of phylogenetic data in the context of human history requires careful assessment of tree topologies, branch lengths and mutation rates as well as thoughtful consilient approaches in integrating historical and archaeological data to prevent overly simplistic, deterministic or even erroneous interpretations.

For the Second Pandemic, the geographic origin and the possible persistence within Europe after the Black Death (1346–1352 AD) are the subject of ongoing scientific and scholarly discussion (Bos et al., 2016; Namouchi et al., 2018; Schmid et al., 2015; Spyrou et al., 2016; Spyrou et al., 2019). Whereas the first comprehensive phylogenetic study on *Y. pestis* favoured an East Asian origin (Cui et al., 2013), other scenarios assume an origin in Central Asia or the Caucasus (Benedictow, 2004; Namouchi et al., 2018; Sussman, 2011). Similarly, the origin of the Justinianic Plague (541–544 AD) has long been hypothesized to have originated in Africa (Achtman et al., 1999; Cui et al., 2008; Sarris, 2002). More recent studies however agree that the strains causing the First Pandemic (541–750 AD) likely emerged in Central Asia (Eroshenko et al., 2017; Harper, 2017; Wagner et al., 2014). The fact that the first outbreak of the Justinianic Plague is reported for Pelusium, Egypt nevertheless raises questions about the history and itinerary of the causative *Y. pestis* strain prior to this outbreak. The currently favoured scenario is an introduction via the Red Sea from India (Harper, 2017; Tsiamis et al., 2009) since there are no historical sources supporting a land route via the Levant or the Arabian peninsula (Schamiloglu, 2016). As such, discrepancies that arise from different analytical approaches raise questions about the history of *Y. pestis* prior to the first documented Justinianic Plague outbreak.

In a recent publication, Damgaard et al., 2018 presented two ancient *Y. pestis* genomes: one from the Tian Shan region (DA101, 2^nd^–3^rd^ c. AD, 8.7-fold average coverage), branching ancestral to the First Pandemic lineage; and one from the Caucasus (DA147, 6^th^–9^th^ c. AD, 0.24-fold average coverage) which was not further investigated. Damgaard et al.’s phylogenetic analysis places DA101 ancestral to the published genome from Aschheim. This positioning is supported by a single SNP shared between the two genomes. An additional five SNPs are unique to DA101 compared to 95 in Aschheim, which is provisionally consistent with Aschheim’s younger age (reported in the SI and in Extended Data Fig. 9, though the latter does not present the tree at full resolution). The identified shared ancestry was interpreted as setting DA101 within the context of the “Justinian plague” (*sic*). The longer branch in the Aschheim genome is explained by its younger age and a seemingly accelerated substitution rate, which is supposedly indicative of an epidemic context.

Although the Justinianic Plague was previously thought to represent the first major onslaught of plague in humans (i.e., the First Pandemic), plentiful examples of human infections of *Y. pestis* are surfacing as far back as the Neolithic (Andrades Valtueña et al., 2017; Rascovan et al., 2019; Spyrou et al., 2018). Here, we present a reanalysis of both DA101 and DA147 genomes which does not seem to support the arguments made by Damgaard et al., 2018. Instead, the analysis of DA101 suggests it to be yet another example of a pre-Justinianic human infection. Furthermore, in contrast to its suggested archaeological dating, we show DA147 to occupy a phylogenetic position much closer to the Black Death (1346–1353 AD) than the Justinianic Plague (Fig. 1C). As such, neither genome can address the origin of the First Pandemic.

**Fig. 1:**
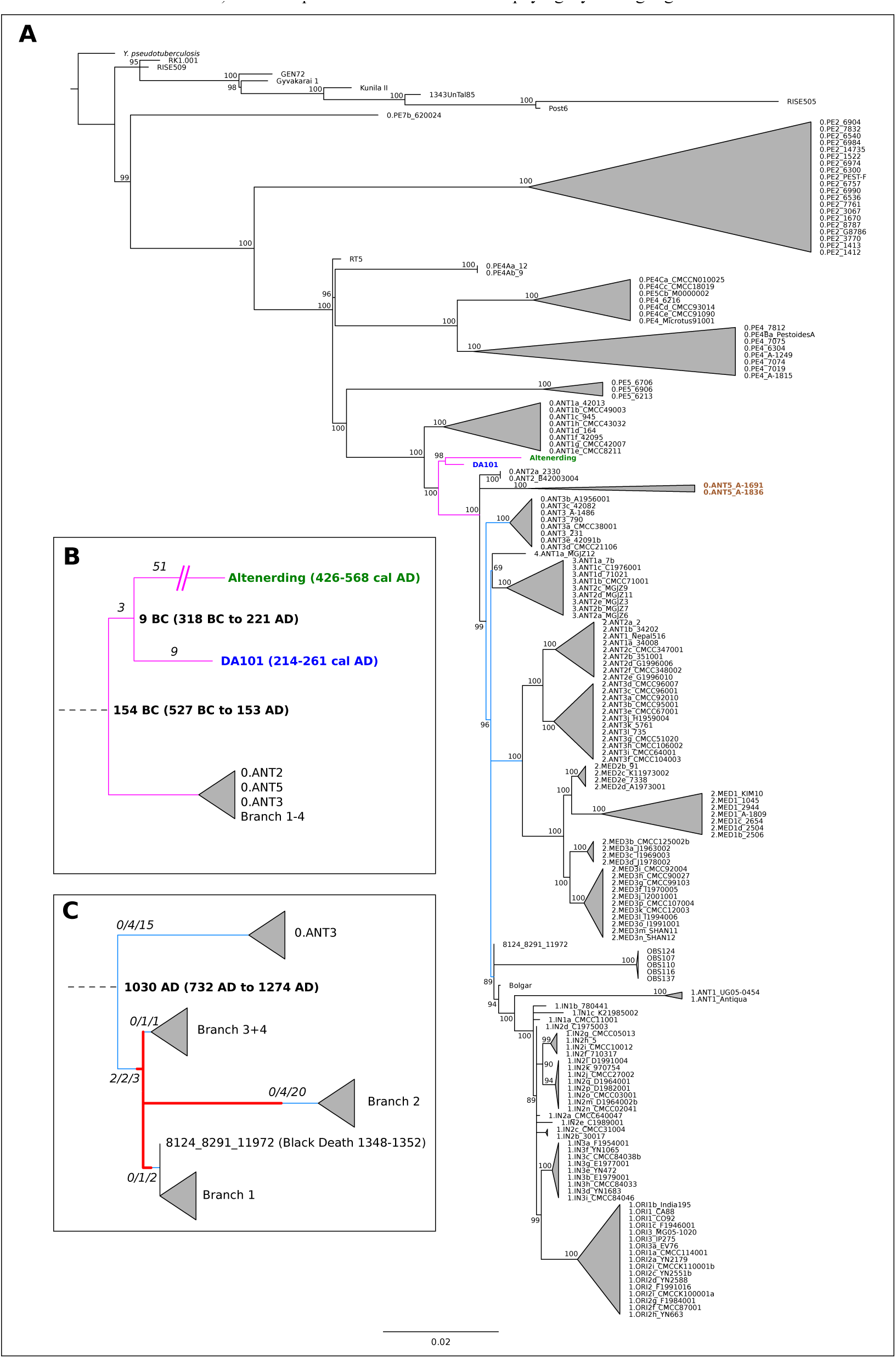
**A.** Maximum likelihood tree based on 3673 SNPs of 167 modern and 18 ancient genomes. Main branches are collapsed for clarity. Numbers on nodes indicate bootstrap support. Highlighted are the Justinianic genome from Altenerding (green), the investigated Tian Shan genome DA101 (blue) and the recently characterized modern strains of clade 0.ANT5 (brown). Relevant parts of the tree are highlighted in pink (Fig. 1B) and light blue (Fig. 1C). **B.** Schematic tree of the common branch of Altenerding and DA101 showing the minimum number of SNPs (italics) and divergence dates (bold, mean and 95 % HPD). **C.** Schematic tree of the highlighted part of A (light blue) for the positioning of DA147 with numbers of SNPs (italics, # of diverged SNPs in DA147/# of covered SNPs in DA147/# of total SNPs on branch) and estimated divergence date of clade 0.ANT3 (bold, mean and 95 % HPD) based on the dating analysis with BEAST 1.10 (see Materials and Methods and SI). Possible placements of DA147 in the phylogeny are highlighted in red.

## Results and Discussion

We reanalysed both presented genomes with a more extensive dataset of published modern and ancient *Y. pestis* genomes (Fig. 1A, Table S1). We opted to include the genome from Altenerding (Feldman et al., 2016) as a representative for the Justinianic Plague: though genetically identical to Aschheim (Wagner et al., 2014), its higher coverage makes it less prone to false positive SNPs that are common in metagenomic data with high environmental backgrounds. Of note, the Aschheim genome has been shown to carry a high number of false positive SNPs (Feldman et al., 2016), which might in part account for its longer branch and accelerated substitution rate observed by Damgaard et al. (see SI).

Analysis of the DA101 genome revealed a minimum of 3 SNPs shared with Altenerding and a minimum of 9 that are unique. By contrast, Altenerding has 51 unique SNPs (Table S2). This sets both nodes, i.e., that giving rise to the shared Justinianic/DA101 branch and the one separating them, deeper in time compared to what is presented in the original publication (Damgaard et al., 2018).

Further to this, we attempted a molecular dating analysis, though the age of individual DA101 proved difficult to determine given discrepancies in the text and SI, ranging from “approximately 180 AD” (main text) to 214–261 calAD/1701 BP (Damgaard et al., 2018 Supplementary Table 2). Ultimately, we opted to use the calibrated radiocarbon interval, which yielded a mean age of 154 BC (95% HPD: 527 BC to 153 AD) for the emergence of the shared lineage and 9 BC (95% HPD: 318 BC to 221 AD) for their divergence time (Table S4). For comparison, dating results without the recently published RT5 genome (Spyrou et al., 2018) are shown in Table S4. This strongly supports a pre-Justinianic provenience for the DA101 genome. A number of shared or unique SNPs might be undetected for DA101 due to low coverage, hence the estimated divergence dates are conservative and might be even older.

Regarding the substitution rate, we do not observe a notable acceleration on the Altenerding branch (mean 2.67E-08) compared to the overall mean (1.48E-08) across the tested dataset (Fig. S2), particularly since both estimates show overlapping 95 % HPD intervals (Table S5). Nevertheless, we do observe an overdispersion of substitution rates across different *Y. pestis* lineages (described previously in Cui et al., 2013 and Spyrou et al., 2019) with the highest estimate here yielding an 17-fold deviation from the mean (2.46E-07).

Damgaard et al. do not discuss the fact that DA101 predates the onset of the Justinianic Plague by three centuries according to its radiocarbon date. This fact, however, is incompatible with their hypothesis of a 6^th^-century pandemic disease introduction to Europe through Hunnic expansion based on this genome alone, as argued suggestively multiple times in their work: in the abstract (“Scythians […] moved westward in about the second or third century BC, forming the Hun traditions in the fourth–fifth century AD, and carrying with them plague that was basal to the Justinian plague.”), the subheader (“Origins and spread of the Justinian plague”, p. 372) and the concluding sentence (“[…], we find provisional support for the hypothesis that the pandemic was brought to Europe towards the end of the Hunnic period through the Silk Road along the southern fringes of the steppes.”, p. 373).

Previously published data demonstrating the absence of detectable genetic changes in *Y. pestis* and its extremely rapid movement during the Black Death in Europe (1347–1353 AD; (Namouchi et al., 2018; Spyrou et al., 2019, 2016) clearly indicate that this pathogen is able to travel vast geographic expanses quickly, without accumulating genetic diversity in the process. As such, the depth of the time interval for the coalescence of DA101 and the Justinianic genomes offers little to no evidence on the temporal or geographic origin of the Justinianic Plague (beginning in 541 AD, Fig. 1B). Since individual DA101 comes from a geographical location that today houses multiple plague foci including modern lineages 0.ANT1, 0.ANT2 and the newly described 0.ANT5 (*sensu* Eroshenko et al., 2017; Fig. 2), it may even be surmised whether the sampled individual fell victim to an epidemic event or a to a sporadic individual infection.

**Fig. 2:**
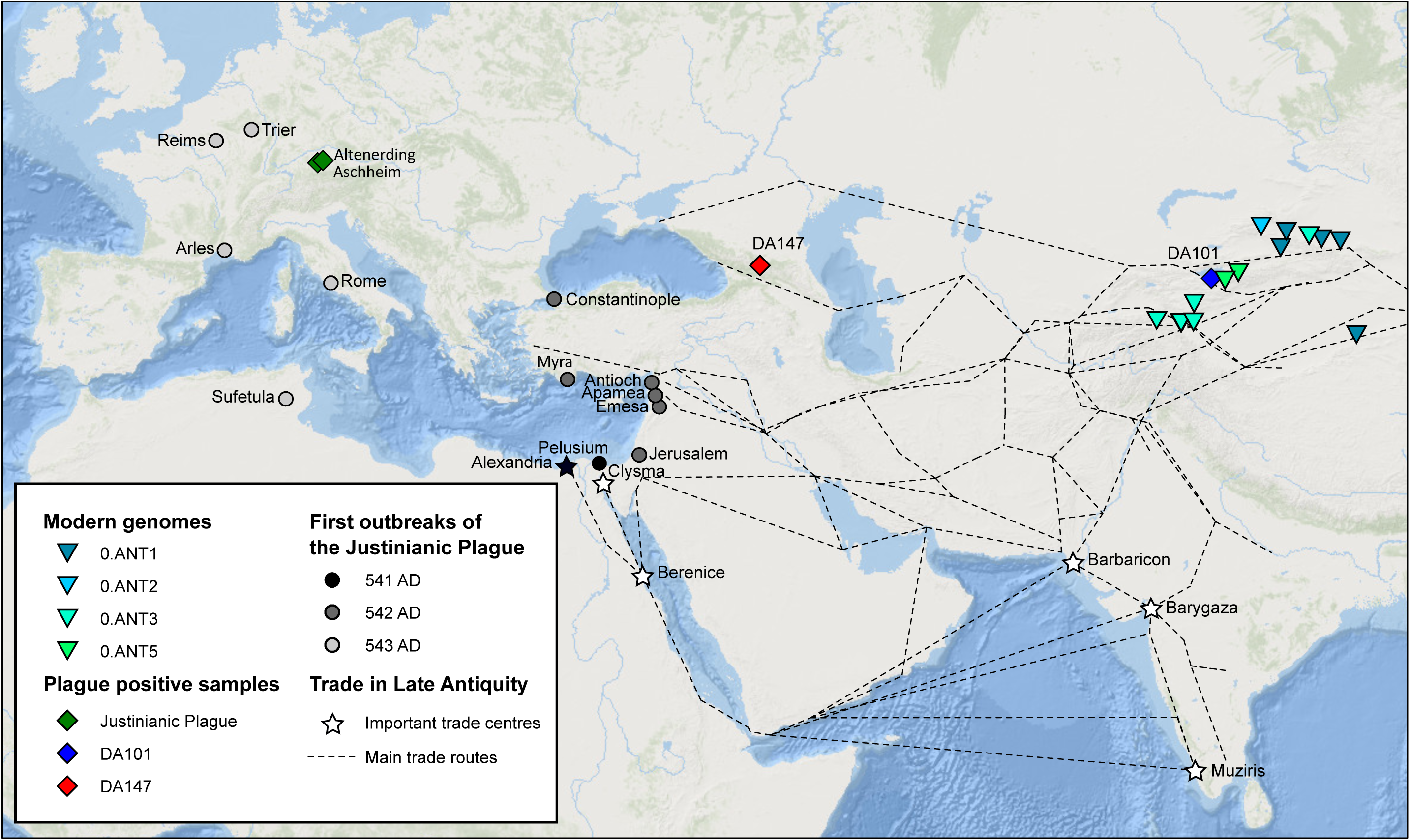
Map of the origin of selected modern and ancient *Y. pestis* genomes in relation to the first outbreaks of the Justinianic Plague as well as important trade centres and routes. An introduction of plague via the Indian Ocean and the Red Sea is supported by well-established sea communications connecting Egyptian and Indian trade centres. The hypothesis of an overland transport through the steppe is in conflict with the directionality of the first outbreaks of the Justinianic Plague and lacks historical support. Although DA147 presumably dates later than 750 AD, it is shown for comparison.

The second *Y. pestis* genome from individual DA147 from North Ossetia, supposedly 6^th^–9^th^ centuries, could substantiate a spread of plague along the “southern fringes of the steppe” (p. 373), although its phylogenetic placement was not investigated by Damgaard et al. Even though the coverage is low, our re-analysis of the raw sequence data from this individual and an assessment of phylogenetically informative positions reveals that it does not share any derived SNPs with Altenerding or DA101 (Table S2). None of the positions shared between Altenerding and DA101 are covered in DA147, but 2 out of the 9 unique SNPs of DA101 are covered and show the ancestral state. Of the unique Altenerding SNPs, 9 are covered in DA147 with 8 showing the ancestral state. The only SNP possibly shared with DA147 is a C>T change that is potentially caused by DNA damage, as it appears only in a single read.

Such initial results motivated a further exploration of DA147’s possible phylogenetic position. For this, we used MultiVCFAnalyzer v0.85 for a comparative SNP analysis against our dataset of ancient and modern *Y. pestis* genomes (Table S1), while omitting all private calls in DA147 since their vast majority will represent DNA damage and sequencing errors due to the genome’s low coverage. The remaining SNPs forming the branch of DA147 in Fig. S1 (red) are an artefact caused by homoplastic or triallelic sites. We computed a maximum likelihood phylogenetic tree that, unexpectedly, placed DA147 closest to the previously described polytomy of Branches 1–4 (Fig. S1). The genomes’s placement was further investigated by visual inspection of all diagnostic SNPs separating Branches 1, 2, 3&4 and Branch 0 (see Table S3). Our analysis reveals several potential placements for DA147: (1) it is one SNP ancestral to the polytomy but derived with respect to the 0.ANT3 node, (2) it is directly on the polytomy, (3) it is one SNP ancestral to the Black Death strain (Bos et al., 2011) on Branch 1, or (4) it is one to 16 SNPs basal on Branch 2 (Fig. 1C; Table S3). The third scenario is of particular interest in the context of a recently discovered genome from Laishevo, Russia (Spyrou et al., 2019) which could be identical to DA147. Therefore, DA147 might instead offer currently unexplored insights into the origin of the Black Death.

Furthermore, this finding raises doubts about the precision in the archaeological dating of this specimen (6^th^–9^th^ centuries; Damgaard et al., 2018). Unfortunately, the provenience of this genome cannot be further investigated since metadata from this individual are absent in Table S2 in Damgaard et al., 2018. Based on our molecular dating analysis, the node giving rise to 0.ANT3, which is basal to all possible placements of DA147, is dated to a mean age of 1030 AD (95% HPD: 732 AD to 1274 AD), thus placing this low coverage genome within the diversity that has accumulated within the last millennium.

Finally, we would like to correct two inaccuracies in nomenclature in the study: First, the label “0.ANT5” has already been given to a modern clade of *Y. pestis* strains reported by Eroshenko et al., 2017. In general, we recommend against applying nomenclature combining phylogenetic and metabolic features to ancient genomes (Achtman, 2016), since their metabolic profile has not yet been characterized. Second, the “Justinianic Plague” is named after the Roman emperor Justinian I (c. 482–565 AD) who reigned during the onset of this pandemic (Little et al., 2007). The term “Justinian Plague” as used by the authors is misleading, since it suggests a connection to either Justin I or Justin II of the Justinianic dynasty.

Overall, we argue that the two presented *Y. pestis* genomes cannot contribute to our understanding of the Justinianic Plague that began in 541 AD in the southeast Mediterranean basin due to their phylogenetic, temporal and geographical distance. Moreover, these genomes offer no support for a connection between the Justinianic Plague and the Hunnic expansion, or for a spread through the southern steppe, both of which are also in conflict with the leading, document-based hypothesis of a plague introduction via trade routes linking India to the Red Sea (Harper, 2017; Fig. 2). The low coverage genome might rather hold clues for the onset of the Black Death or on the origins of Branch 2. We suggest a redirected focus here, especially if higher coverage data from this or a similar archaeological sample becomes available in the future.

## Materials and Methods

Sequencing data for the samples DA101 and DA147 were retrieved from ENA with the provided accession numbers (Damgaard et al., 2018) and processed with the EAGER pipeline (Peltzer et al., 2016), including Illumina adapter removal, sequencing quality filtering (minimum base quality of 20) and length filtering (minimum length of 30 bp).

For the DA101 sample with higher coverage, reads were clipped on both ends by 3 bases to remove the majority of damaged sites and subsequently filtered again for length using the same parameter. Mapping against the CO92 reference genome (chromosome NC_003143.1) was done with BWA (-l 32, -n 0.1, -q 37), reads with low mapping quality (-q 37) were removed with Samtools and duplicates were removed with MarkDuplicates.

SNP calling was performed with the UnifiedGenotyper within the Genome Analysis Toolkit (GATK) using the ‘EMIT_ALL_SITES’ option to generate a call for every position in the reference genome.

For the DA147 sample with low coverage, mapping was performed without prior damage clipping and with less stringent parameters in BWA (-l 16, -n 0.01, -q 37) to retrieve a maximum of coverage. Reads with low mapping quality were removed with Samtools (-q 37) and duplicates were removed with MarkDuplicates. For a phylogenetic analysis of the low coverage DA147 genome (0.24-fold), the bam-file was converted into a fastq-file using bedtools, multiplied by 5 and mapped again with identical parameters but without duplicate removal to reach the necessary coverage of positions for SNP calling. SNP calling was performed with the UnifiedGenotyper within the Genome Analysis Toolkit using ‘EMIT_ALL_SITES’ to generate calls for all positions in the reference genome.

For the phylogenetic analyses, we used 166 previously published modern *Y. pestis* genomes (Cui et al., 2013; Eroshenko et al., 2017; Kislichkina et al., 2015; Zhgenti et al., 2015), a *Y. pseudotuberculosis* reference genome (IP32953; Chain et al., 2004) as an outgroup and the following ancient genomes: nine genomes from Neolithic/Bronze Age contexts (Andrades Valtueña et al., 2017; Rasmussen et al., 2015; Spyrou et al., 2018), one genome of the Justinianic Plague (Altenerding; Feldman et al., 2016), one genome representing Black Death (8291-11972-8124; (Bos et al., 2011), and six genomes of the subsequent second plague pandemic (Observance OBS116, OBS137, OBS110, OBS107, OBS124; Bos et al., 2016); Bolgar, (Spyrou et al., 2016)). A complete list of all *Y. pestis* genomes used is given in Table S1. Previously identified problematic regions as well as regions annotated as repeat regions, rRNAs, tRNAs and tmRNAs were excluded for all subsequent analyses (Cui et al., 2013; Morelli et al., 2010). MultiVCFAnalyzer v0.85 (Bos et al., 2014) was used for generating a SNP table with the following settings: Minimal coverage for base call of 5 with a minimum genotyping quality of 30 for homozygous positions, minimum support of 90% for calling the dominant nucleotide in a ‘heterozygous’ position. The sample DA147 was processed in the outgroup mode in MultiVCFAnalyzer to remove all singletons, for the most part representing damaged sites called due to the prior multiplication of reads. The unfiltered SNP alignment produced by MultiVCFAnalyzer was used for all following analyses. Additionally, all phylogenetically informative positions were visually inspected in IGV v2.4 (Thorvaldsdóttir et al., 2013). Maximum likelihood trees were generated with RAxML v8 (Stamatakis, 2014) using the GTR substitution model based on a partial deletion (95 %) SNP alignment for DA101 (3673 SNPs) and a full SNP alignment for DA147 (3885 SNPs). Robustness of all trees was tested by the bootstrap methods using 1000 pseudo-replicates.

For the estimation of divergence times and substitution rates with BEAST 1.10 (Drummond and Rambaut, 2007), we used the coalescent Bayesian skyline model with a setup identical to that published in Spyrou et al., 2018 with the following modifications: Integration of the DA101 sample (Damgaard et al., 2018), 95% partial deletion SNP alignment and 800,000,000 states as chain length. A second run was performed without the recently published Bronze Age genome RT5 (Spyrou et al., 2018) to investigate whether the results are affected by previously unavailable data. The recently published genomes by Eroshenko et al., 2017 were not included in the dating analysis due to exceptionally long branches and unavailability of raw data to address potential mismapping. An MCC tree was produced using TreeAnnotator of BEAST v1.10, showing the relative mean substitution rates (Fig. S2). All trees were visualized in FigTree v1.4.3 (http://tree.bio.ed.ac.uk/software/figtree/).

For Fig. 2, we used the coordinates of DA146 and DA160, since they are identical and frame the sample DA147 which is not part of the Table S2 of Damgaard et al., 2018. The first outbreaks of the Justinianic Plague come from Harper, 2017 and (Stathakopoulos, 2004). For the trade routes, we used the “Indian and Persian trade routes with the West 50 BCE - 300 CE” and “Silk Road routes 1–1400 CE” from OWTRAD (http://www.ciolek.com/owtrad.html). Important trade centres are adopted from Harper, 2017.

## Author Contributions

M.K., A.H., K.I.B and J.K. planned and designed the study. M.K. performed data processing and phylogenetic analyses; M.A.S. and M.K. performed dating analyses. M.M. provided and reviewed historical context information. M.K. wrote the manuscript with contributions from M.A.S., K.I.B. and A.H. and edits from all co-authors.

