## Supplementary Information for "Ancient *Yersinia pestis* genomes provide no evidence for the origins or spread of the Justinianic Plague"

**Fig. S1:** Maximum likelihood tree based on 3885 SNPs of 167 modern and 19 ancient genomes. Main branches are collapsed for clarity, numbers on nodes indicate bootstrap support. Highlighted are the Justinianic genome from Altenerding (green), the investigated Tian Shan genome DA101 (blue). DA147 and its artificial branch are shown in red, all branches relevant for the positioning of DA147 are shown in light blue (see Fig. 1A and C).

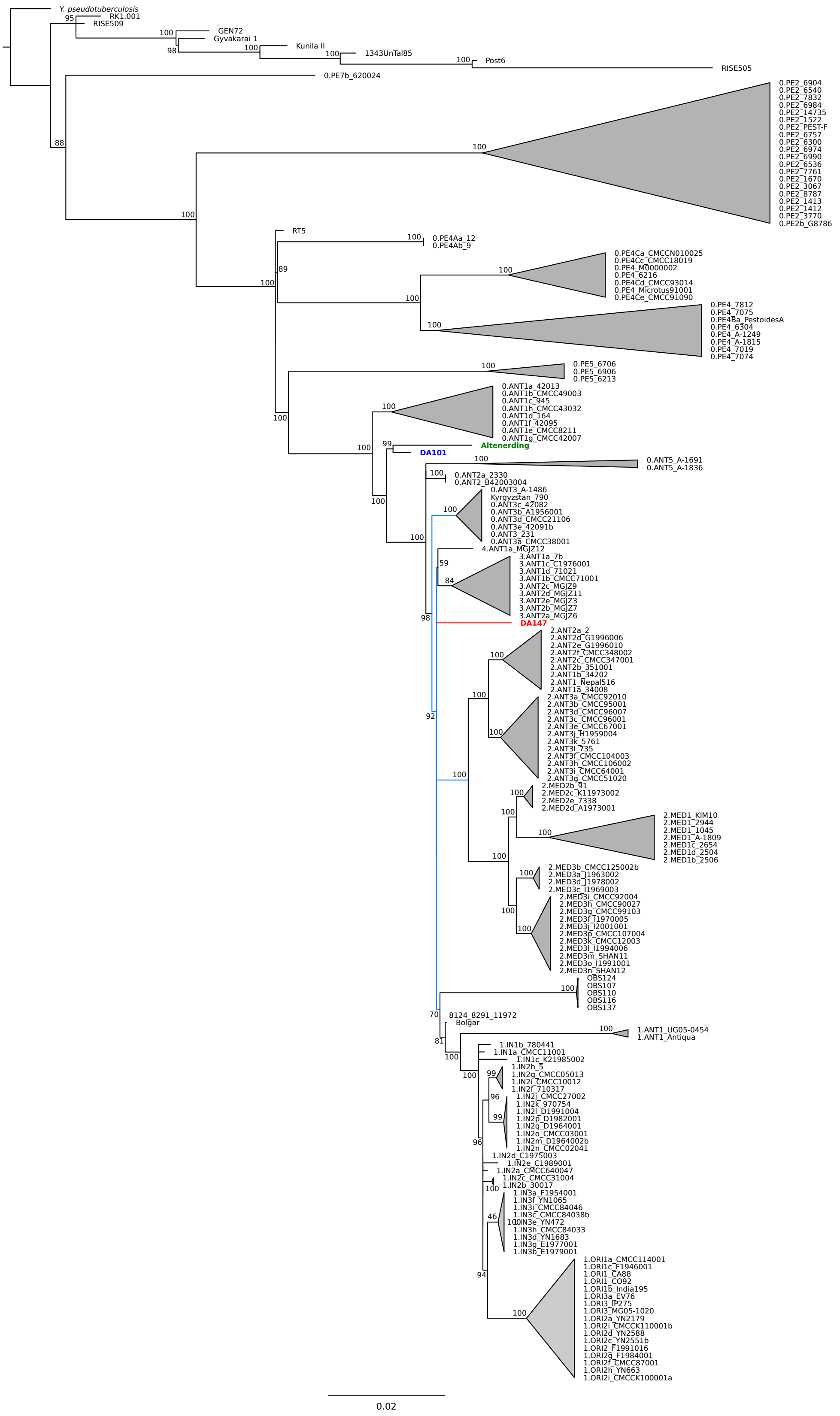

**Figure S2:** Maximum clade credibility tree showing substitution rate across all branches of the *Y. pestis* phylogeny, years BP are indicated on the bottom scale. Since the analysis is based on a SNP alignment with partial deletion (95%).

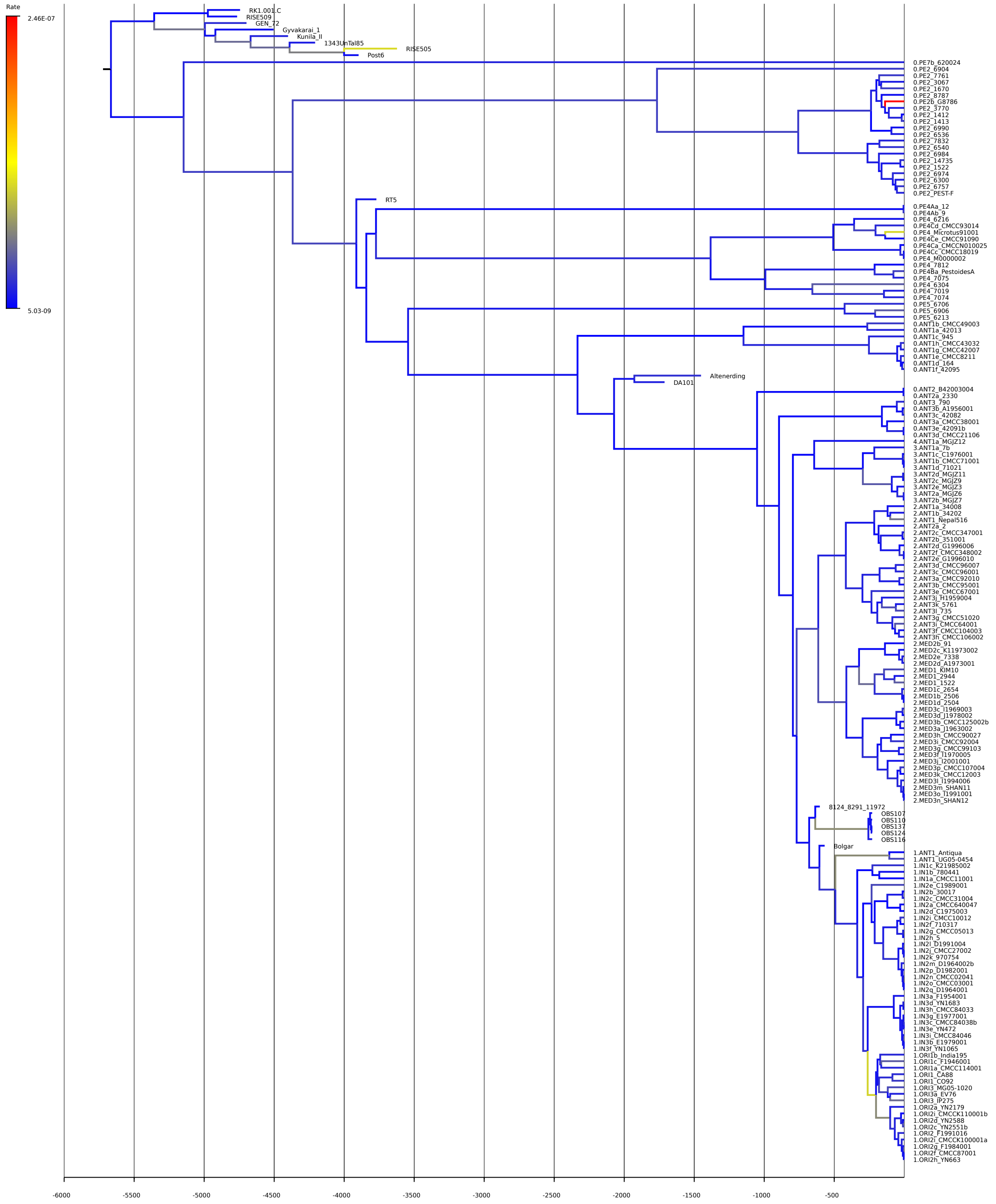

**Table S1:** List of all *Y. pestis* genomes with accession number, sample origin and the corresponding publication.

| Strain ID | Accession No. | Origin | Publication |
|---|---|---|---|
| **Modern** | | | |
| 0.ANT1a_42013 | ADPG00000000 | Xinjiang, China | Cui et al. 2013 |
| 0.ANT1b_CMCC49003 | ADQX00000000 | Xinjiang, China | Cui et al. 2013 |
| 0.ANT1c_945 | ADPV00000000 | Xinjiang, China | Cui et al. 2013 |
| 0.ANT1d_164 | ADOW00000000 | Xinjiang, China | Cui et al. 2013 |
| 0.ANT1e_CMCC8211 | ADRD00000000 | Xinjiang, China | Cui et al. 2013 |
| 0.ANT1f_42095 | ADPJ00000000 | Xinjiang, China | Cui et al. 2013 |
| 0.ANT1g_CMCC42007 | ADQV00000000 | Xinjiang, China | Cui et al. 2013 |
| 0.ANT1h_CMCC43032 | ADQW00000000 | Xinjiang, China | Cui et al. 2013 |
| 0.ANT2_B42003004 | AAYU00000000 | Xinjiang, China | Cui et al. 2013 |
| 0.ANT2a_2330 | ADOY00000000 | Xinjiang, China | Cui et al. 2013 |
| 0.ANT3_231 | JMUF00000000 | Kyrgyzstan | Eroshenko et al. 2017 |
| 0.ANT3_790 | CP006806 | Kyrgyzstan | Zhgenti et al. 2015 |
| 0.ANT3_A_1496 | LYMP00000000 | Kyrgyzstan | Eroshenko et al. 2017 |
| 0.ANT3a_CMCC38001 | ADQU00000000 | Xinjiang, China | Cui et al. 2013 |
| 0.ANT3b_A1956001 | ADPX00000000 | Xinjiang, China | Cui et al. 2013 |
| 0.ANT3c_42082 | ADPH00000000 | Xinjiang, China | Cui et al. 2013 |
| 0.ANT3d_CMCC21106 | ADQP00000000 | Xinjiang, China | Cui et al. 2013 |
| 0.ANT3e_42091b | ADPI00000000 | Xinjiang, China | Cui et al. 2013 |
| 0.ANT5_A_1691 | LYMQ00000000 | Kyrgyzstan | Eroshenko et al. 2017 |
| 0.ANT5_A_1836 | LYOL00000000 | Kyrgyzstan | Eroshenko et al. 2017 |
| 0.PE2_14735 | AYLS00000000 | Armenia | Zhgenti et al. 2015 |
| 0.PE2_1522 | CP006758 | Armenia | Zhgenti et al. 2015 |
| 0.PE2_1412 | CP006783 | Dalidag, Georgia | Zhgenti et al. 2015 |
| 0.PE2_1413 | CP006762 | Ninotsminda, Georgia | Zhgenti et al. 2015 |
| 0.PE2_1670 | AYLR00000000 | Ninotsminda, Georgia | Zhgenti et al. 2015 |
| 0.PE2_3067 | CP006754 | Akhalkalaki, Georgia | Zhgenti et al. 2015 |
| 0.PE2_3770 | CP006751 | Ninotsminda, Georgia | Zhgenti et al. 2015 |
| 0.PE2_8787 | CP006748 | Ninotsminda, Georgia | Zhgenti et al. 2015 |
| 0.PE2_6300 | LIZC00000000 | Pre-Araks, Armenia, Azerbaijan | Kislichkina et al. 2015 |
| 0.PE2_6536 | LIZE00000000 | Transcaucasian Highland, Armenia, Georgia | Kislichkina et al. 2015 |
| 0.PE2_6540 | LIZF00000000 | Transcaucasian Highland, Armenia, Azerbaijan | Kislichkina et al. 2015 |
| 0.PE2_6757 | LIYY00000000 | Pre-Araks, Armenia, Azerbaijan | Kislichkina et al. 2015 |
| 0.PE2_6904 | LIYP00000000 | Dagestan-highland, Russia | Kislichkina et al. 2015 |
| 0.PE2_6974 | LIYX00000000 | Transcaucasian Highland, Armenia, Azerbaijan | Kislichkina et al. 2015 |
| 0.PE2_6984 | LIYZ00000000 | Transcaucasian Highland, Armenia, Azerbaijan | Kislichkina et al. 2015 |
| 0.PE2_6990 | LIYU00000000 | Transcaucasian Highland, Armenia, Georgia | Kislichkina et al. 2015 |
| 0.PE2_7761 | LIYQ00000000 | Transcaucasian Highland, Armenia, Georgia | Kislichkina et al. 2015 |
| 0.PE2_7832 | LIZB00000000 | Transcaucasian Highland, Armenia, Azerbaijan | Kislichkina et al. 2015 |
| 0.PE2_PEST-F | NC_009381 | Former Soviet Union | Cui et al. 2013 |
| 0.PE2b_G8786 | ADSG00000000 | Georgia | Cui et al. 2013 |
| 0.PE4_6216 | LIYR00000000 | Bayan-Khongor, Mongolia | Kislichkina et al. 2015 |
| 0.PE4_6304 | LIYS00000000 | Gissar, Tadjikistan, Uzbekistan | Kislichkina et al. 2015 |
| 0.PE4_7019 | LIYW00000000 | Talas, Kyrgyzstan | Kislichkina et al. 2015 |
| 0.PE4_7074 | LIYT00000000 | Talas, Kyrgyzstan | Kislichkina et al. 2015 |
| 0.PE4_7075 | LIZA00000000 | Mountain-Altai, Russia | Kislichkina et al. 2015 |
| 0.PE4_7812 | LIYV00000000 | Mountain-Altai, Russia | Kislichkina et al. 2015 |
| 0.PE4_Microtus91001 | NC_005810 | Inner Mongolia, China | Cui et al. 2013 |
| 0.PE4Aa_12 | ADOV00000000 | Qinghai, China | Cui et al. 2013 |
| 0.PE4Ab_9 | ADPT00000000 | Qinghai, China | Cui et al. 2013 |
| 0.PE4b_M0000002 | ADST00000000 | Qinghai, China | Cui et al. 2013 |
| 0.PE4Ba_PestoidesA | ACNT00000000 | Former Soviet Union | Cui et al. 2013 |
| 0.PE4Ca_CMCCN010025 | ADRT00000000 | Sichuan, China | Cui et al. 2013 |
| 0.PE4Cc_CMCC18019 | ADQO00000000 | Qinghai, China | Cui et al. 2013 |
| 0.PE4Cd_CMCC93014 | ADRM00000000 | Inner Mongolia, China | Cui et al. 2013 |
| 0.PE4Ce_CMCC91090 | ADRJ00000000 | Inner Mongolia, China | Cui et al. 2013 |
| 0.PE4h_A_1249 | LYMN00000000 | Tadzhikistan | Eroshenko et al. 2017 |
| 0.PE4t_A_1815 | LPTY00000000 | Kyrgyzstan | Eroshenko et al. 2017 |
| 0.PE5_6213 | LIZD00000000 | Northeast Mongolia, Gobi Desert | Kislichkina et al. 2015 |
| 0.PE5_6706 | LIYO00000000 | Northeast Mongolia, Gobi Desert | Kislichkina et al. 2015 |
| 0.PE5_6906 | LIZG00000000 | Northeast Mongolia, Gobi Desert | Kislichkina et al. 2015 |
| 0.PE7b_620024 | ADPM00000000 | Qinghai, China | Cui et al. 2013 |
| 1.ANT1_Antiqua | NC_008150 | Congo | Cui et al. 2013 |
| 1.ANT1_UG05-0454 | AAYR00000000 | Uganda | Cui et al. 2013 |
| 1.IN1a_CMCC11001 | ADQK00000000 | Qinghai, China | Cui et al. 2013 |
| 1.IN1b_780441 | ADPS00000000 | Qinghai, China | Cui et al. 2013 |

| | | | |
|---|---|---|---|
| 1.IN1c_K21985002 | ADSS00000000 | Xinjiang, China | Cui et al. 2013 |
| 1.IN2a_CMCC640047 | ADRA00000000 | Qinghai, China | Cui et al. 2013 |
| 1.IN2b_30017 | ADPC00000000 | Tibet, China | Cui et al. 2013 |
| 1.IN2c_CMCC31004 | ADQR00000000 | Tibet, China | Cui et al. 2013 |
| 1.IN2d_C1975003 | ADPZ00000000 | Qinghai, China | Cui et al. 2013 |
| 1.IN2e_C1989001 | ADQB00000000 | Qinghai, China | Cui et al. 2013 |
| 1.IN2f_710317 | ADPP00000000 | Qinghai, China | Cui et al. 2013 |
| 1.IN2g_CMCC05013 | ADQF00000000 | Qinghai, China | Cui et al. 2013 |
| 1.IN2h_5 | ADPK00000000 | Qinghai, China | Cui et al. 2013 |
| 1.IN2i_CMCC10012 | ADQG00000000 | Qinghai, China | Cui et al. 2013 |
| 1.IN2j_CMCC27002 | ADQQ00000000 | Qinghai, China | Cui et al. 2013 |
| 1.IN2k_970754 | ADPW00000000 | Qinghai, China | Cui et al. 2013 |
| 1.IN2l_D1991004 | ADRX00000000 | Qinghai, China | Cui et al. 2013 |
| 1.IN2m_D1964002b | ADRV00000000 | Qinghai, China | Cui et al. 2013 |
| 1.IN2n_CMCC02041 | ADQC00000000 | Qinghai, China | Cui et al. 2013 |
| 1.IN2o_CMCC03001 | ADQD00000000 | Qinghai, China | Cui et al. 2013 |
| 1.IN2p_D1982001 | ADRW00000000 | Gansu, China | Cui et al. 2013 |
| 1.IN2q_D1964001 | ADRU00000000 | Qinghai, China | Cui et al. 2013 |
| 1.IN3a_F1954001 | ADSC00000000 | Yunnan, China | Cui et al. 2013 |
| 1.IN3b_E1979001 | AAYV00000000 | Yunnan, China | Cui et al. 2013 |
| 1.IN3c_CMCC84038b | ADRF00000000 | Yunnan, China | Cui et al. 2013 |
| 1.IN3d_YN1683 | ADTD00000000 | Yunnan, China | Cui et al. 2013 |
| 1.IN3e_YN472 | ADTH00000000 | Yunnan, China | Cui et al. 2013 |
| 1.IN3f_YN1065 | ADTC00000000 | Yunnan, China | Cui et al. 2013 |
| 1.IN3g_E1977001 | ADRY00000000 | Yunnan, China | Cui et al. 2013 |
| 1.IN3h_CMCC84033 | ADRE00000000 | Yunnan, China | Cui et al. 2013 |
| 1.IN3i_CMCC84046 | ADRG00000000 | Yunnan, China | Cui et al. 2013 |
| 1.ORI1_CA88 | ABCD00000000 | California, USA | Cui et al. 2013 |
| 1.ORI1_CO92 | NC_003143 | Colorado, USA | Cui et al. 2013 |
| 1.ORI1a_CMCC114001 | ADQL00000000 | Fujian, China | Cui et al. 2013 |
| 1.ORI1b_India195 | ACNR00000000 | India | Cui et al. 2013 |
| 1.ORI1c_F1946001 | ADSB00000000 | Fujian, China | Cui et al. 2013 |
| 1.ORI2_F1991016 | ABAT00000000 | Yunnan, China | Cui et al. 2013 |
| 1.ORI2a_YN2179 | ADTE00000000 | Myanmar | Cui et al. 2013 |
| 1.ORI2c_YN2551b | ADTF00000000 | Yunnan, China | Cui et al. 2013 |
| 1.ORI2d_YN2588 | ADTG00000000 | Guangxi, China | Cui et al. 2013 |
| 1.ORI2f_CMCC87001 | ADRH00000000 | Yunnan, China | Cui et al. 2013 |
| 1.ORI2g_F1984001 | ADSD00000000 | Yunnan, China | Cui et al. 2013 |
| 1.ORI2h_YN663 | ADTI00000000 | Yunnan, China | Cui et al. 2013 |
| 1.ORI2i_CMCCK100001a | ADRR00000000 | Yunnan, China | Cui et al. 2013 |
| 1.ORI2i_CMCCK110001b | ADRS00000000 | Yunnan, China | Cui et al. 2013 |
| 1.ORI3_IP275 | AAOS00000000 | Madagascar | Cui et al. 2013 |
| 1.ORI3_MG05-1020 | AAYS00000000 | Madagascar | Cui et al. 2013 |
| 1.ORI3a_EV76 | ADSA00000000 | Madagascar | Cui et al. 2013 |
| 2.ANT1_Nepal516 | ACNQ00000000 | Nepal | Cui et al. 2013 |
| 2.ANT1a_34008 | ADPD00000000 | Tibet, China | Cui et al. 2013 |
| 2.ANT1b_34202 | ADPE00000000 | Tibet, China | Cui et al. 2013 |
| 2.ANT2a_2 | ADOX00000000 | Qinghai, China | Cui et al. 2013 |
| 2.ANT2b_351001 | ADPF00000000 | Tibet, China | Cui et al. 2013 |
| 2.ANT2c_CMCC347001 | ADQS00000000 | Tibet, China | Cui et al. 2013 |
| 2.ANT2d_G1996006 | ADSE00000000 | Tibet, China | Cui et al. 2013 |
| 2.ANT2e_G1996010 | ADSF00000000 | Tibet, China | Cui et al. 2013 |
| 2.ANT2f_CMCC348002 | ADQT00000000 | Tibet, China | Cui et al. 2013 |
| 2.ANT3a_CMCC92010 | ADRL00000000 | Inner Mongolia, China | Cui et al. 2013 |
| 2.ANT3b_CMCC95001 | ADRN00000000 | Inner Mongolia, China | Cui et al. 2013 |
| 2.ANT3c_CMCC96001 | ADRO00000000 | Inner Mongolia, China | Cui et al. 2013 |
| 2.ANT3d_CMCC96007 | ADRP00000000 | Inner Mongolia, China | Cui et al. 2013 |
| 2.ANT3e_CMCC67001 | ADRB00000000 | Inner Mongolia, China | Cui et al. 2013 |
| 2.ANT3f_CMCC104003 | ADQH00000000 | Inner Mongolia, China | Cui et al. 2013 |
| 2.ANT3g_CMCC51020 | ADQY00000000 | Jilin, China | Cui et al. 2013 |
| 2.ANT3h_CMCC106002 | ADQI00000000 | Inner Mongolia, China | Cui et al. 2013 |
| 2.ANT3i_CMCC64001 | ADQZ00000000 | Inner Mongolia, China | Cui et al. 2013 |
| 2.ANT3j_H1959004 | ADSI00000000 | Jilin, China | Cui et al. 2013 |
| 2.ANT3k_5761 | ADPL00000000 | St.Petersburg, Russia | Cui et al. 2013 |
| 2.ANT3l_735 | ADPR00000000 | St.Petersburg, Russia | Cui et al. 2013 |
| 2.MED1_A_1809 | LYMF00000000 | Kyrgyzstan | Eroshenko et al. 2017 |
| 2.MED1_1045 | CP006794 | Azerbaijan | Zhgenti et al. 2015 |
| 2.MED1_2944 | CP006792 | Kabardino-Balkaria, Russia | Zhgenti et al. 2015 |
| 2.MED1b_2506 | ADPA00000000 | Xinjiang, China | Cui et al. 2013 |
| 2.MED1c_2654 | ADPB00000000 | Xinjiang, China | Cui et al. 2013 |
| 2.MED1d_2504 | ADOZ00000000 | Xinjiang, China | Cui et al. 2013 |

| | | | |
|---|---|---|---|
| 2.MED2_KIM10 | NC_004088 | Iran/Kurdistan | Cui et al. 2013 |
| 2.MED2b_91 | ADPU00000000 | Xinjiang, China | Cui et al. 2013 |
| 2.MED2c_K11973002 | AAYT00000000 | Xinjiang, China | Cui et al. 2013 |
| 2.MED2d_A1973001 | ADPY00000000 | Xinjiang, China | Cui et al. 2013 |
| 2.MED2e_7338 | ADPQ00000000 | Xinjiang, China | Cui et al. 2013 |
| 2.MED3a_J1963002 | ADSP00000000 | Gansu, China | Cui et al. 2013 |
| 2.MED3b_CMCC125002b | ADQN00000000 | Ningxia, China | Cui et al. 2013 |
| 2.MED3c_I1969003 | ADSK00000000 | Ningxia, China | Cui et al. 2013 |
| 2.MED3d_J1978002 | ADSQ00000000 | Ningxia, China | Cui et al. 2013 |
| 2.MED3f_I1970005 | ADSL00000000 | Inner Mongolia, China | Cui et al. 2013 |
| 2.MED3g_CMCC99103 | ADRQ00000000 | Inner Mongolia, China | Cui et al. 2013 |
| 2.MED3h_CMCC90027 | ADRI00000000 | Inner Mongolia, China | Cui et al. 2013 |
| 2.MED3i_CMCC92004 | ADRK00000000 | Inner Mongolia, China | Cui et al. 2013 |
| 2.MED3j_I2001001 | ADSO00000000 | Shaanxi, China | Cui et al. 2013 |
| 2.MED3k_CMCC12003 | ADQM00000000 | Qinghai, China | Cui et al. 2013 |
| 2.MED3l_I1994006 | ADSN00000000 | Hebei, China | Cui et al. 2013 |
| 2.MED3m_SHAN11 | ADTA00000000 | Shaanxi, China | Cui et al. 2013 |
| 2.MED3n_SHAN12 | ADTB00000000 | Shaanxi, China | Cui et al. 2013 |
| 2.MED3o_I1991001 | ADSM00000000 | Inner Mongolia, China | Cui et al. 2013 |
| 2.MED3p_CMCC107004 | ADQJ00000000 | Inner Mongolia, China | Cui et al. 2013 |
| 3.ANT1a_7b | ADPN00000000 | Qinghai, China | Cui et al. 2013 |
| 3.ANT1b_CMCC71001 | ADRC00000000 | Gansu, China | Cui et al. 2013 |
| 3.ANT1c_C1976001 | ADQA00000000 | Gansu, China | Cui et al. 2013 |
| 3.ANT1d_71021 | ADPO00000000 | Gansu, China | Cui et al. 2013 |
| 3.ANT2a_MGJZ6 | ADSX00000000 | Dornogovi, Mongolia | Cui et al. 2013 |
| 3.ANT2b_MGJZ7 | ADSY00000000 | Dornogovi, Mongolia | Cui et al. 2013 |
| 3.ANT2c_MGJZ9 | ADSZ00000000 | Govi-Altai, Mongolia | Cui et al. 2013 |
| 3.ANT2d_MGJZ11 | ADSU00000000 | Bayan-Ölgii, Mongolia | Cui et al. 2013 |
| 3.ANT2e_MGJZ3 | ADSW00000000 | Govi-Altai, Mongolia | Cui et al. 2013 |
| 4.ANT1a_MGJZ12 | ADSV00000000 | Bayan-Ölgii, Mongolia | Cui et al. 2013 |
| **Ancient** | | | |
| 8124_8291_11972 | SRR341961,<br>SRR341962,<br>SRR341963 | United Kingdom | Bos et al. 2011 |
| RISE505 | PRJEB10885 | Russia | Rasmussen et al. 2015 |
| RISE509 | PRJEB10885 | Russia | Rasmussen et al. 2015 |
| OBS107 | PRJEB12163 | France | Bos et al. 2016 |
| OBS110 | PRJEB12163 | France | Bos et al. 2016 |
| OBS116 | PRJEB12163 | France | Bos et al. 2016 |
| OBS124 | PRJEB12163 | France | Bos et al. 2016 |
| Altenerding | PRJEB14851 | Germany | Feldman et al. 2016 |
| Bolgar | PRJEB13664 | Russia | Spyrou et al. 2016 |
| GEN72 | PRJEB19335 | Croatia | Andrades Valtueña et al. 2017 |
| RK1.001.C | PRJEB19335 | Russia | Andrades Valtueña et al. 2017 |
| Kunila II | PRJEB19335 | Estonia | Andrades Valtueña et al. 2017 |
| Gyvakarai 1 | PRJEB19335 | Lithuania | Andrades Valtueña et al. 2017 |
| 6Post | PRJEB19335 | Germany | Andrades Valtueña et al. 2017 |
| 1343UnTal85 | PRJEB19335 | Germany | Andrades Valtueña et al. 2017 |
| DA101 | PRJEB25891 | Kyrgyzstan | Daamgard et al. 2018 |
| DA147 | PRJEB25891 | Russia | Daamgard et al. 2018 |
| RT5 | PRJEB24296 | Russia | Spyrou et al. 2018 |

**Table S2:** Unique and shared SNPs of DA101 and Altenerding. *Erroneously classified as ancestral in Feldman et al. 2016. **Potential damage site (C>T).

| **Shared SNPs** | | | | |
|---|---|---|---|---|
| **Position** | **Reference (CO92)** | **Altenerding** | **DA101** | **DA147** |
| 260148 | C | T | T | N |
| 2725715 | C | T | T | N |
| 2977542 | C | A | A | N |
| **Unique SNPs Altenerding** | | | | |
| **Position** | **Reference (CO92)** | **Altenerding** | **DA101** | **DA147** |
| 86824 | A | G | N | N |
| 189912 | A | G | . | N |
| 420208* | G | T | . | . |
| 271114 | C | A | . | N |
| 485976 | C | T | . | . |
| 557841 | C | T | N | N |
| 727741 | G | A | . | N |
| 779365 | C | T | . | N |
| 898980 | A | T | N | . |
| 1067966 | C | A | . | N |
| 1211729 | A | C | . | N |
| 1296743 | C | T | . | N |
| 1387701 | C | T | N | N |
| 1387756 | A | G | . | . |
| 1413031 | C | A | N | . |
| 1434752 | C | A | . | N |
| 1489055 | C | T | . | N |
| 1530658 | C | A | . | N |
| 1609461 | T | C | N | N |
| 1754708 | C | T | . | T* |
| 1868678 | G | T | N | N |
| 1956162 | T | C | . | N |
| 2092152 | C | T | . | N |
| 2097520 | G | T | . | N |
| 2352174 | T | G | . | . |
| 2419529 | G | A | . | N |
| 2495165 | C | A | . | N |
| 2753572 | A | T | . | N |
| 3078807 | C | A | . | N |
| 3179828 | C | A | . | N |
| 3274298 | A | T | N | N |
| 3360963 | A | C | . | N |
| 3360984 | C | T | . | N |
| 3398153 | G | A | N | N |
| 3409414 | T | C | N | N |
| 3500922 | T | G | . | N |
| 3535148 | G | T | . | N |
| 3560088 | G | A | . | . |
| 3568597 | C | T | . | N |
| 3750736 | G | A | N | N |
| 3755861 | C | T | N | . |
| 3843195 | C | A | . | N |
| 3892488 | C | T | . | N |
| 4066494 | C | T | N | N |
| 4307755 | G | A | . | N |
| 4412624 | A | G | . | . |
| 4423366 | G | A | N | N |

| | | | | |
|---|---|---|---|---|
| 4460688 | C | T | N | N |
| 4465967 | C | A | N | N |
| 4628496 | C | A | . | N |
| 4629169 | G | A | N | N |
| **Unique SNPs DA101** | | | | |
| **Position** | **Reference (CO92)** | **Altenerding** | **DA101** | **DA147** |
| 100945 | C | . | T | N |
| 178109 | C | . | T | N |
| 3066176 | C | . | T | N |
| 3075453 | G | . | T | . |
| 3550137 | C | . | T | N |
| 3592088 | C | . | A | N |
| 3641873 | G | . | A | N |
| 3686087 | C | . | T | N |
| 3729628 | G | . | A | . |

**Table S3:** SNPs defining possible positions of DA147 in the phylogenetic tree.

| **Position** | **Reference (CO92)** | **0.ANT3** | **DA147** | **State** |
|---|---|---|---|---|
| 561567 | G | A | N | not covered |
| 1440879 | A | G | N | not covered |
| 1698435 | A | G | N | not covered |
| 1719187 | C | T | N | not covered |
| 2003542 | C | T | N | not covered |
| 2082310 | C | T | N | not covered |
| 2130133 | G | T | N | not covered |
| 2425991 | G | T | . | ancestral |
| 2656734 | C | T | . | ancestral |
| 3512754 | T | C | . | ancestral |
| 3727189 | G | T | N | not covered |
| 4166664 | G | A | . | ancestral |
| 4245783 | C | T | N | not covered |
| 4281601 | G | T | N | not covered |
| 4427796 | T | G | N | not covered |
| **Position** | **Reference (CO92)** | **Branch 0** | **DA147** | **State** |
| 1102174 | A | G | . | derived |
| 1251046 | T | C | N | not covered |
| 2812384 | G | T | . | derived |
| **Position** | **Reference (CO92)** | **Branch 1** | **DA147** | **State** |
| 189227 | C | . | T | ancestral |
| 1871476 | G | . | N | not covered |
| **Position** | **Reference (CO92)** | **Branch 2** | **DA147** | **State** |
| 97226 | G | A | . | ancestral |
| 282762 | G | A | N | not covered |
| 335833 | C | T | N | not covered |
| 443673 | G | T | N | not covered |
| 710909 | C | T | N | not covered |
| 718827 | T | C | N | not covered |
| 759797 | G | T | . | ancestral |
| 1971665 | G | A | N | not covered |
| 2082100 | C | T | N | not covered |
| 2493895 | C | T | N | not covered |
| 2799652 | T | C | N | not covered |
| 2847692 | C | T | N | not covered |
| 2934864 | C | T | N | not covered |
| 3314013 | G | T | N | not covered |
| 3376387 | A | G | . | ancestral |
| 3539709 | C | T | N | not covered |
| 3551089 | A | C | N | not covered |
| 4311918 | C | T | N | not covered |
| 4595001 | C | T | . | ancestral |
| 4598609 | G | A | N | not covered |
| **Position** | **Reference (CO92)** | **Branch 3+4** | **DA147** | **State** |
| 3021936 | C | A | . | ancestral |

**Table S4:** Mean divergence dates and 95 % HPD intervals for important nodes in the phylogenetic tree.

| Lineage divergence | Mean (95 % HPD) in years BP including RT5 | Mean (95 % HPD) in years BP excluding RT5 |
|---|---|---|
| Tree root | 5782 (4941–6863) | 5449 (4869–6200) |
| 0.PE7 | 5232 (4229–6672) | 4490 (3235–5912) |
| 0.PE2 | 4428 (3904–5073) | 3483 (1642–4453) |
| 0.PE4 (incl. RT5) | 3932 (3742–4153) | 2803 (2175–3492) |
| 0.PE5 | 3478 (2763–4004) | 2605 (2058–3235) |
| 0.ANT1 | 2372 (1924–2891) | 2139 (1846–2476) |
| Altenerding+DA101 | 2104 (1797–2477) | 1977 (1770–2244) |
| DA101 | 1959 (1729–2268) | 1881 (1722–2091) |
| 0.ANT2 | 1085 (741–1503) | 986 (703–1348) |
| 0.ANT3 | 920 (676–1218) | 844 (652–1101) |
| Black Death | 654 (602–763) | 637 (602–712) |

**Table S5:** Mean substitution rates and their respective 95% HPD for the whole Y. pestis phylogeny and the Altenerding branch.

| | Mean (95 % HPD) including RT5 | Mean (95 % HPD) excluding RT5 |
|---|---|---|
| Full tree | 1.48E-08 (1.23E-08–1.73E-08) | 1.75E-08 (1.42E-08–2.08E-08) |
| Altenerding | 2.67E-08 (1.08E-08–4.36E-08) | 3.08E-08 (1.08E-08–4.82E-08) |
